# A distinct phylogenetic cluster of Indian SARS-CoV-2 isolates

**DOI:** 10.1101/2020.05.31.126136

**Authors:** Sofia Banu, Bani Jolly, Payel Mukherjee, Priya Singh, Shagufta Khan, Lamuk Zaveri, Sakshi Shambhavi, Namami Gaur, Rakesh K Mishra, Vinod Scaria, Divya Tej Sowpati

## Abstract

From an isolated epidemic, COVID-19 has now emerged as a global pandemic. The availability of genomes in the public domain following the epidemic provides a unique opportunity to understand the evolution and spread of the SARS-CoV-2 virus across the globe. The availability of whole genomes from multiple states in India prompted us to analyse the phylogenetic clusters of genomes in India. We performed whole-genome sequencing for 64 genomes making a total of 361 genomes from India, followed by phylogenetic clustering, substitution analysis, and dating of the different phylogenetic clusters of viral genomes. We describe a distinct phylogenetic cluster (Clade I / A3i) of SARS-CoV-2 genomes from India, which encompasses 41% of all genomes sequenced and deposited in the public domain from multiple states in India. Globally 3.5% of genomes, which till date could not be mapped to any distinct known cluster fall in this newly defined clade. The cluster is characterized by a core set of shared genetic variants – C6312A (T2016K), C13730T (A88V/A97V), C23929T, and C28311T (P13L). Further, the cluster is also characterized by a nucleotide substitution rate of 1.4 × 10^−3^ variants per site per year, lower than the prevalent A2a cluster, and predominantly driven by variants in the E and N genes and relative sparing of the S gene. Epidemiological assessments suggest that the common ancestor emerged in the month of February 2020 and possibly resulted in an outbreak followed by countrywide spread, as evidenced by the low divergence of the genomes from across the country. To the best of our knowledge, this is the first comprehensive study characterizing the distinct and predominant cluster of SARS-CoV-2 in India.

## Introduction

Since the emergence of the outbreak in the Chinese city of Wuhan in late 2019, the novel Coronavirus disease has spread widely to become a global pandemic, with over 5,000,000 individuals infected worldwide and resulting in the death of over 300,000 individuals (as of 26 May 2020).^1^ The causative virus, SARS-CoV-2, is a member of the genus betacoronavirus. During its transmission, the virus has differentiated into at least 10 clades globally and is continuously evolving.^2^ This has implications in genetic epidemiology, surveillance, contact tracing and the development of long term strategies for mitigation of this disease.^3^

The recent availability of whole-genome sequences of the SARS-CoV-2 from across the world deposited in public databases provides an unprecedented opportunity to understand the dynamics and evolution of the pathogen. The availability of genomic data in a public repository like GISAID^4^ also provides wider access to the resources and enables researchers across the globe to address pertinent hypotheses. Likewise, this gave us a unique scope to understand the introduction, evolution, and spread of the virus in India and understand it in the context of global clades circulating across the world.

In this manuscript, we report the sequences of SARS-CoV-2 isolates predominantly sampled from the state of Telangana. Further, we systematically analysed the phylogenetic clusters of genomes from India and characterised a unique cluster of sequences (Clade I/A3i), which could not be classified into any of the previously annotated global clades. Isolates forming this cluster were predominant in a number of states and potentially characterized by a shared set of four genetic variants. The cluster potentially arose from a single outbreak followed by a rapid spread across the country. To the best of our knowledge, this is the first comprehensive report of the novel and predominant cluster of sequences from India and suggests its distribution beyond India in many countries in the Middle East, South Asia, and Oceania.

## Materials and Methods

### Sample collection and RNA purification

Samples were collected and processed as per the guidelines of the Institutional Ethics Committee. Nasopharyngeal or oropharyngeal swabs collected in viral transport media were used for RNA isolation. Samples were anonymised by removing all patient identifiers except for gender, age, collection date, and symptoms where applicable. SARS-CoV-2 nucleic acids were isolated from 300 µl viral transport media using the QIAamp Viral RNA Mini kit (Qiagen) according to the manufacturer’s protocol. Samples were eluted in 50 µl nuclease-free water and stored in −80°C until further use.

### Shotgun RNA sequencing

100ng of RNA was taken as input for rRNA depletion using TruSeq RiboZero Gold kit (Illumina). Following rRNA depletion and fragmentation, RNA was converted to cDNA and processed for NGS library prep using TruSeq Stranded Total RNA library prep kit (Illumina). Library profiles were analyzed using Bioanalyzer and sequenced on Illumina NovaSeq 6000 at a depth of 30 million reads per sample.

### Targeted viral genome sequencing

A modified version of the nCoV-2019 sequencing protocol as described by Josh Quick, 2020 was used to amplify the viral genome.^5^ Briefly, a combination of random hexamers, anchored oligo(dT)23, and the 98R primer from nCoV-2019/V3 primer set were used to synthesize the first strand according to the reaction conditions mentioned in the protocol. Next, a 3-step multiplex PCR was carried out to amplify the viral genome using nCoV-2019/V3 primer pools 1 and 2. The ∼400 bp amplicons thus obtained in two pools were combined, purified using Agencourt AMPure XP beads (Beckman Coulter) and eluted in 30 µl elution buffer (Qiagen). Around 100 ng of purified amplicons were used for library preparation. DNA libraries were prepared using either the QIAseq FX DNA Library Kit (Qiagen) or Nextera DNA Flex Library Prep Kit (Illumina) according to the manufacturer’s protocol. Paired-End Sequencing (2×150bp) was performed on the NovaSeq 6000 (Illumina) with a targeted depth of 2 million reads per sample (∼20,000X coverage).

### Assembly of sequencing data

Raw image data was basecalled using bcl2fastq v2.20 with --no-lane-splitting option enabled. QC of the FASTQ files was performed using FastQC v0.11.7, and adaptors/poor quality bases were trimmed using Trimmomatic.^6,7^ Reads were aligned to the reference genome MN908947.3 using hisat2.^8^ Consensus sequence from the bam file was derived using seqtk and bcftools.^9^ Samtools depth command was used to calculate the coverage across the genome, with the options -a and -d 50000.^10^ The sequences were deposited in GISAID. The complete list of accessions is detailed in **Supplementary Data 1**.

### Genomic Data and Analysis

The datasets of Indian SARS-CoV-2 genomes deposited in GISAID (till 25 May 2020) were used for the analysis. The samples, sample annotations, originating, and submitting institutions are listed in **Supplementary Data 2**. Further 10 high-quality genomes from each of the 10 clades respectively as annotated by Nextstrain were retrieved from GISAID and used in the analysis.

### Quality filters

A stringent quality criteria was employed for considering the genomes for evaluation of the nucleotide substitution rates, molecular clock, and phylogenetic clustering, as these would be sensitive to the quality of genomes. For this, the individual samples were systematically aligned to the Wuhan-Hu-1 reference genome (NC_045512) using EMBOSS needle.^11^ Samples which had Ns > 1% and/or gaps > 1% of the sequence length and/or degenerate bases were removed from the analysis. Similarly, if a submitting laboratory had >90% samples annotated as low quality, all the data from the laboratory was removed from the analysis. Samples having an ambiguous date of collection were also removed from the analysis. After the initial high-quality analysis, all genomes originating from India irrespective of the quality criteria were considered for computation of the proportions of clades in each of the states.

### Masking of low-quality variants

A compendium of problematic genomic loci was compiled from a previous study and from the protocol specified by Nextstrain.^12,13^ These positions were masked in all the genomes. This compendium included 36 variants and is available as **Supplementary Data 3**.

### Phylogenetic clustering of genomes

Phylogenetic analysis of the samples was performed as detailed previously following the standard protocol for analysis of SARS-CoV-2 genomes provided by Nextstrain.^13,14^ The sequences were clustered using Augur, the phylodynamic pipeline provided by Nextstrain.^15^ The sequences were aligned against the WH1 reference genome using MAFFT and spurious variant positions were masked from the alignment.^16^ The initial phylogenetic tree was constructed using IQTREE following the Augur tree implementation.^17^ The raw tree was further processed with Augur to construct a TimeTree, annotate ancestral traits, infer mutations, and identify clades. The resulting tree was viewed using Nextstrain.

### Divergence estimates and most recent common ancestor

The substitution rate of nCoV-2 was estimated using BEAST v1.10.4 following a methodology described previously.^18,19^ The masked alignment file was used as input to estimate a strict molecular clock and a coalescent growth rate model (exponential growth). MCMC was run for 50 million steps and burn-in was adjusted to attain a suitable effective sample size (ESS).

Times to the most recent common ancestor (tMRCA) for the individual clusters were also computed by Bayesian coalescent analysis using BEAST. All runs were executed using the HKY+Г substitution model with gamma-distributed rate variation (gamma categories=4), a strict molecular clock and a coalescent growth rate model. MCMC was run for steps as described above. The log output of each MCMC run was analysed in Tracer v1.7.1.^20^ The priors used for each parameter in the analysis are given in **Supplementary Data 4**.

### Functional evaluation of variants

The variants were also evaluated for the functional consequences using the Sorting Intolerant from Tolerant (SIFT).^21^ SIFT prediction database for SARS-CoV-2 was built using the genome annotations for the WH1 reference genome (NC_045512). A SIFT score of 0.0 to 0.05 was interpreted to have a deleterious effect while scores in the range 0.05 to 1.0 were predicted to be tolerated. The functional effects of protein variants identified in the clades were assessed using the PROVEAN web server, using the protein sequences of the Wuhan-Hu-1 genome as reference and a default threshold value of −2.5.^22^ Additionally, PhyloP conservation scores and base-wise GERP Rejected Substitutions scores (RS scores) for the variants were also computed.^23,24^ Sites having positive PhyloP scores are predicted to be conserved, while negative scores are indicative of fast-evolving sites. Positive GERP scores were considered indicative of a site under evolutionary constraint, while negative scores indicate neutrally evolving sites. The variants were also checked for overlaps with immune epitope predictions as given on UCSC Genome Browser for SARS-CoV-2.

## Results

### Demographics and quality of viral genomes

The samples sequenced encompass 64 genomes in total, majorly collected from the state of Telangana. The age of the patients ranged from 10-73 years, with >80% (54 out of 64) within the age bracket of 20-60 years (Fig S1A). A total of 61 samples were sequenced using an amplicon-based approach with a target of ∼2 million paired-end reads per sample. We could achieve an average coverage of >1000x in all cases, with a uniform representation from all amplicons (Fig S1B top, S1C). 3 samples were sequenced using a shotgun sequencing approach and had an average coverage of approximately 100x, however, the coverage across the genome was uniform (Fig S1B, bottom). The samples and metadata for the isolates sequenced and deposited in the public domain are summarised in **Supplementary Data 1**

### Phylogenetic Clusters

A total of 361 genome sequences of the SARS-CoV-2 were available for analysis as of 25th May 2020 from India including the genomes sequenced by our group. After stringent analysis of the sequence quality and removal of all sequences which did not meet the quality criteria, the dataset resulted in a total of 213 genomes submitted from 7 institutions (including 55 out of our 64 genomes). **Supplementary Data** 2 summarises the genomic data considered for the analysis, submitting institutions, quality criteria used for inclusion/exclusion, and the acknowledgements.

The genomes isolated from India were found to be classified under 5 clusters. 4 of these clusters are known clades identified by Nextstrain: A2a, A3, B, and B4.^13^The first and the major cluster encompassed 133 (62%) of genomes which fell into the A2a clade of the SARS-CoV-2 genome. The clade was represented by samples derived from multiple states across the country including Gujarat, West Bengal, Maharashtra, Tamil Nadu, and Telangana.

The second-largest cluster consisted of 62 genomes (29%). This cluster of sequences could not be classified into any of the 10 clade sequences defined by Nextstrain.^13^ We further evaluated the nucleotide compositions which define each of the 10 clusters/clades worldwide and compared this to the nucleotide composition of the clade in question. Our analysis suggests that the nucleotide compositions which define the 10 clades were absent in the clade of question. This cluster was found to have diverged from the A1a and A3 clades and most, but not all of the sequences shared a variant (L3606F in ORF1a) with members of the A3 and A1a clades. We propose to call this the A3i clade in cognisance of this fact. To avoid potential conflict with the nomenclature followed by Nextstrain, we, therefore, define this cluster of sequences as Clade I/A3i, for the unique occurrence as a dominant cluster amongst SARS-CoV-2 genome sequences from India and also since this clade is largely formed by sequences from India.

The other clusters encompassed the A3, B4, and B clades with 13, 3, and 1 genomes each falling into the respective clusters. Figure 1 summarises the different phylogenetic clusters of the genomes from India.

**Figure 1.**
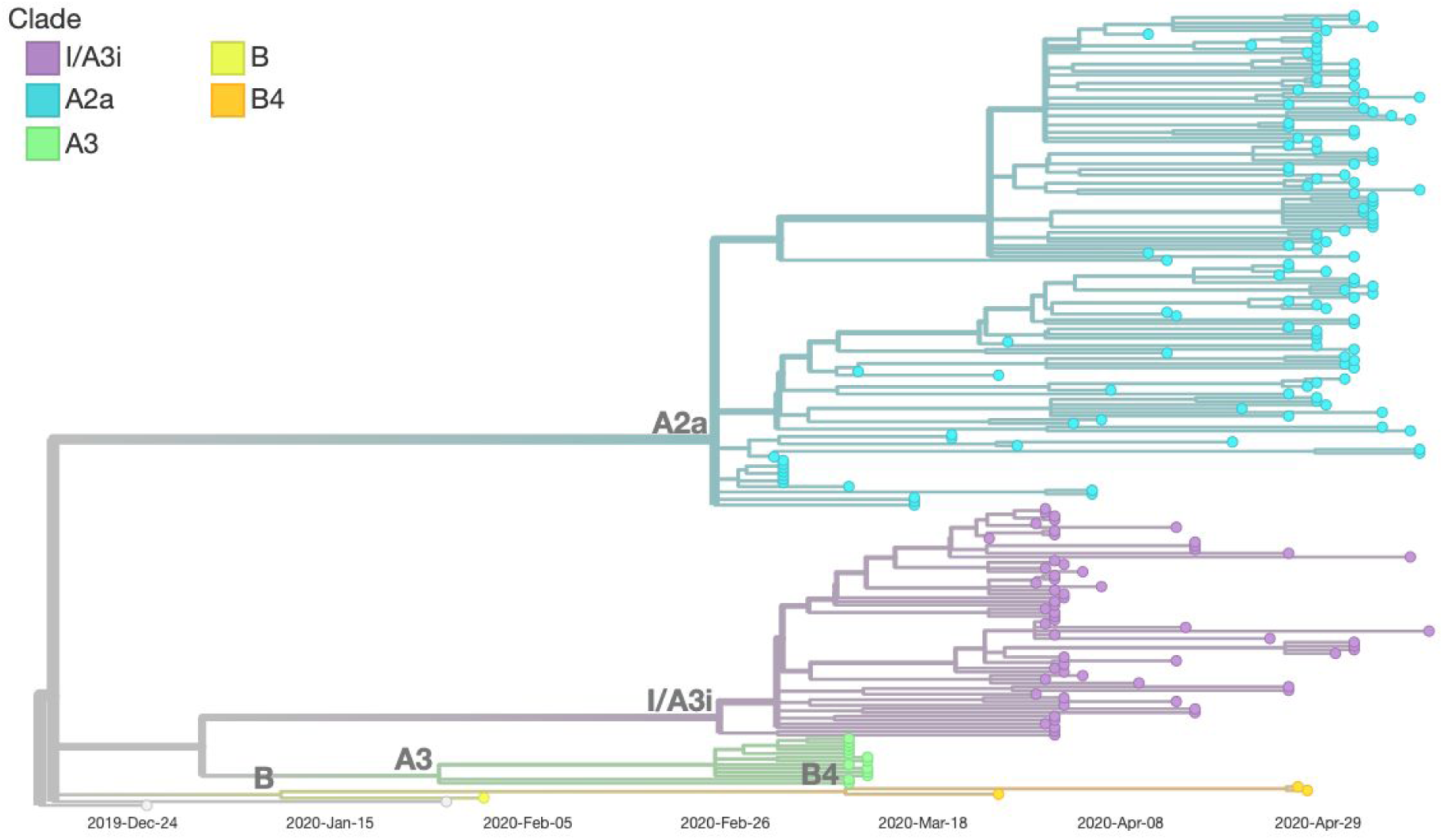
Phylogenetic clusters and the clades assigned for the Indian SARS-CoV-2 genomes.

### Molecular definition of the cluster

Systematic analysis of members of the cluster revealed that a set of unique genetic variants defined the core cluster. A discriminant analysis was performed for all variants in any genome defined by the cluster of sequences. A unique combination of four variants, C6312A, C13730T, C23929T, and C28311T was found to be shared by a majority of members of the cluster. A total of 60 genomes of the 62 genomes (96.8%) in the cluster shared the combination of variants. This unique combination of variants was shared by none of the other genomes which were assigned to any other clade.

We further analysed the global datasets for identifying the genomes which displayed matches for 4/4 variants which defined the Clade I/A3i. Our prospective search retrieved a total of 21 high-quality genomes (**Supplementary Data 5**). Of the retrieved genomes, the largest number of genomes originated from Singapore which had 5 genomes and constituted 8% of the high-quality genomes from Singapore. The other genomes originated from a number of countries including Australia, Brazil, Brunei, Canada, China, Gambia, Guam, Philippines, Saudi Arabia, Slovenia, and Taiwan. The members in the cluster, however, contributed to a much smaller proportion of the clades/clusters identified in the respective countries.

Of these, 4 were sampled from a date earlier than the earliest sample of this cluster from India and were from Australia, Brazil, Canada, and Saudi Arabia.

### Nucleotide Substitution Rates

Mutation rates were calculated for the Indian sequences using BEAST, with the WH1 genome as the root. Our analysis suggests that the substitution rate is 1.64 × 10^−3^ (95% HPD 1.41 × 10^−3^ – 1.88 × 10^−3^) substitutions per site per year for the entire Indian SARS-CoV-2 genomes put together. This also confirms the estimates previously made.^25^

The substitution rate was also computed for the individual clades. The gene-wise substitution rates were also similarly calculated for the major clusters. The analysis suggests that the I/A3i clade has a nucleotide substitution rate of 1.4 × 10^−3^ variants per site per year compared to the estimate of 1.65 × 10^−3^ variants per site per year for the prevalent A2a clade and 1.64 × 10^−3^ variants per site per year for all the high quality genomes from India analysed.

The nucleotide substitution rate suggests that the evolution of the I/A3i clade is largely determined by changes in the structural proteins – Nucleocapsid (N) and Envelope (E) genes, compared to the A2a, the globally predominant clade, which is determined by changes in the Spike (S) and Membrane (M) genes (Table 1).

**Table 1.** Nucleotide Substitution rates of the different structural protein genes and genome-wide across the different clusters and clades in India. The estimates for B and B4 were not computed since the clades had only 3 and 1 genomes respectively from India.

| Clade/<br>Cluster | S Gene | E Gene | M Gene | N Gene | Genome |
| --- | --- | --- | --- | --- | --- |
| **All (N=213)** | $3.10 \times 10^{-3}$ | $1.89 \times 10^{-3}$ | $3.6 \times 10^{-3}$ | $4.5 \times 10^{-3}$ | $1.64 \times 10^{-3}$ |
| **A2a (N=133)** | $3.15 \times 10^{-3}$ | $0.79 \times 10^{-3}$ | $2.57 \times 10^{-3}$ | $1.78 \times 10^{-3}$ | $1.65 \times 10^{-3}$ |
| **I/A3i (N=62)** | $1.13 \times 10^{-3}$ | $2.96 \times 10^{-3}$ | $2.37 \times 10^{-3}$ | $3.52 \times 10^{-3}$ | $1.4 \times 10^{-3}$ |
| **A3 (N=13)** | $1.04 \times 10^{-3}$ | $1.98 \times 10^{-3}$ | $0.63 \times 10^{-3}$ | $5.48 \times 10^{-3}$ | $1.55 \times 10^{-3}$ |

### Estimating time to most recent common ancestor and age of the cluster

The date of the most recent ancestor for the dataset of all Indian SARS-CoV-2 genomes, with WH1 genome sequence included, was computed using BEAST. The median tMRCA was found to be 11 December 2019 (95% HPD 26 November to 25 December), confirming the previous estimates of the origin of the epidemic in Wuhan city of China.^26^

tMRCA for the I/A3i clade, as well as the A2a clade, which constituted the majority of samples, was also computed. The clade A2a, which is the predominant clade in India had a tMRCA of 2nd Jan 2020 (95% HPD Interval 13 Dec 2019 – 22 Jan 2020), while the clade I/A3i had a tMRCA of 8 Feb 2020 (95% HPD Interval 17 Jan 2020 – 25 Feb 2020).

### Functional consequences of the variants

Of the four variants which characterise the clade I/A3i, three of the variants caused an amino acid change (non-synonymous). Additionally, all variants which defined the other clades were compiled. The nonsynonymous variants were analysed for potential functional consequences using the Sorting Intolerant from Tolerant (SIFT) and Protein Variant Effect Analyser (PROVEAN).^21,22^ SIFT prediction database for SARS-CoV-2 was built using the genome annotations for the WH1 reference genome (NC_045512). Additionally, PhyloP and GERP conservation scores were also computed for the variants.^23,24^ The variants and their respective predictions are summarised in **Table 2**.

**Table 2.** Functional characteristics of the four variants which define Clade I/A3i and other clades across the world. PROVEAN scores of less than −2.5 are considered deleterious in nature. Similarly, SIFT scores of 0 to 0.05 are considered deleterious.

| Clade | Gene | Site | Mutation | PROVEAN score / Prediction | SIFT score / Prediction | Conservation Scores |
| --- | --- | --- | --- | --- | --- | --- |
| A1a | ORF3a | G26144T | G251V | -8.581<br>Deleterious | 0<br>Deleterious | PhyloP: 4.256<br>GERP: 1.65 |
| A1a, A3, I/A3i | ORF1a | G11083T | L3606F | -1.4<br>Neutral | 0.01<br>Deleterious | PhyloP: -1.32286<br>GERP: -3.3 |
| A2 | S | A23403G | D614G | 0.598<br>Neutral | 0.3<br>Tolerated | PhyloP: 2.25839<br>GERP: 1.65 |
| A2a | ORF1b | C14408T | P314L | -0.914<br>Neutral | 0.31<br>Tolerated | PhyloP: 3.30748<br>GERP: 1.65 |
| A3 | ORF1a | G1397A | V378I | -0.199<br>Neutral | 0.62<br>Tolerated | PhyloP: 0.227575<br>GERP: -1.81 |
| A7 | ORF1a | C9924T | A3220V | -2.049<br>Neutral | 0.04<br>Deleterious | PhyloP: 3.30935<br>GERP: 1.65 |
| B, B1, B2, B4 | ORF8 | T28144C | L84S | 2.333<br>Neutral | 0.37<br>Tolerated | PhyloP: -1.52089<br>GERP: -0.206 |
| B4 | N | G28878A | S202N | -0.404<br>Neutral | 0<br>Deleterious | PhyloP: 4.256<br>GERP: 1.65 |
| I/A3i | N | C28311T | P13L | -1.23<br>Neutral | 0<br>Deleterious | PhyloP: 3.27687<br>GERP: 1.65 |
| I/A3i | RDRP /ORF1b | C13730T | A97V (A88V in ORF1b) | -3.611<br>Deleterious | 0<br>Deleterious | PhyloP: 3.31844<br>GERP: 1.65 |
| I/A3i | ORF1a | C6312A | T2016K | -0.352<br>Neutral | 0.03<br>Deleterious | PhyloP: 3.29661<br>GERP: 1.61 |

Majority of the variants which defined other clades were predicted to be neutral by PROVEAN, with the exception of G251V which defines the A1a clade. Three variants which define Clade I/A3i, C6312A, C13730T and C28311T resulted in amino acid changes with potentially deleterious functional consequences as predicted by SIFT and mapped to quite conserved genomic loci in the SARS-CoV-2 genome **(Table 2)**. One of these variants, A97V in the RDRP protein (corresponds to A88V in ORF1b) is located in its NiRAN domain, which is suggested to be important in RNA binding and nucleotidylation activity.^27^ Both SIFT and PROVEAN analyses **(Table 2)** suggest that the effect of this mutation is deleterious in nature, however, as both Alanine and Valine are hydrophobic amino acids, the exact effect of the mutation needs to be experimentally validated. Of notable significance is the P13L variant (C28311T) in the Nucleocapsid protein which is required for the viral entry into the cells. The variant maps to the IDR domain of the N protein and SIFT predicts the variant to be deleterious, though the PROVEAN analysis categorized it as a neutral mutation **(Table 2)**.

Two of the variants C6312A in ORF1a and C13730T in ORF1b also mapped to immune epitope predictions (HLA-A0201 binding peptides) from NetMHC 4.0 available on UCSC Genome Browser and as listed on UCSC Genome Browser for SARS-CoV-2.^28^ The potential consequences of the variants in the immune response could not be ascertained.

### Defining the origin and spread from the cluster

The presence of a short tree of Clade I/A3i with divergence from a single point suggests a single point of introduction.^29^ The single point of divergence also suggests that the origin and spread of the cluster were possibly from a single outbreak **(Figure 3)**. The clustering of samples in Feb 2020 suggests a rapid spread spanning multiple regions across the country. The first sequence from the cluster in India was GMC-KN443/2020 (Accession ID EPI_ISL_431103, deposited by Department of Microbiology, Gandhi Medical College and Hospital, Hyderabad, India) sampled on 16th of March from an Indonesian traveller from the state of Telangana. Of the 6 states from which the data for high-quality genomes were made available, the I/A3i clade was represented in 4 of the states.

**Figure 2.**
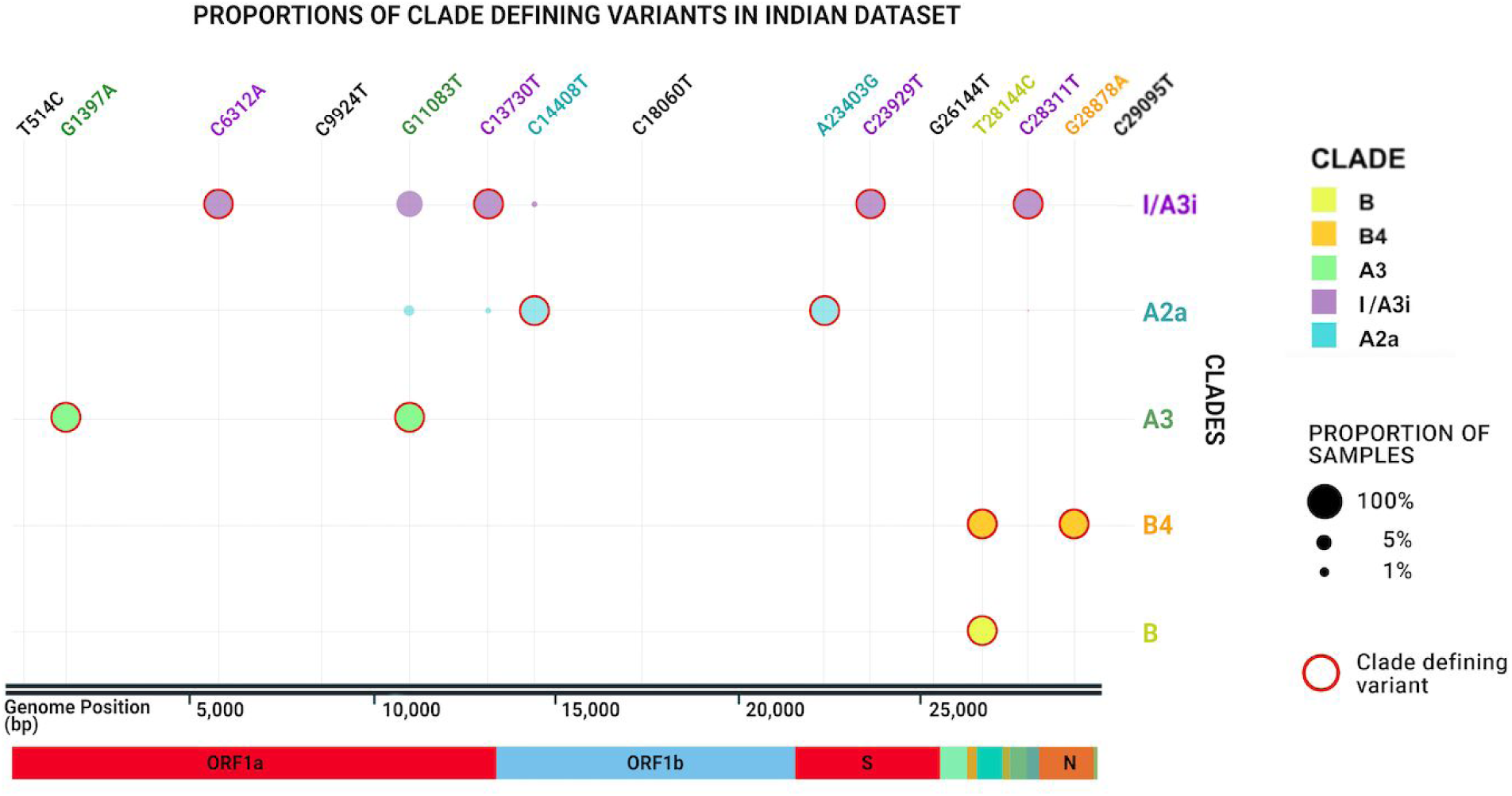
Shared variants among the variants which define the clusters of the Indian SARS-CoV-2 genomes.

**Figure 3.**
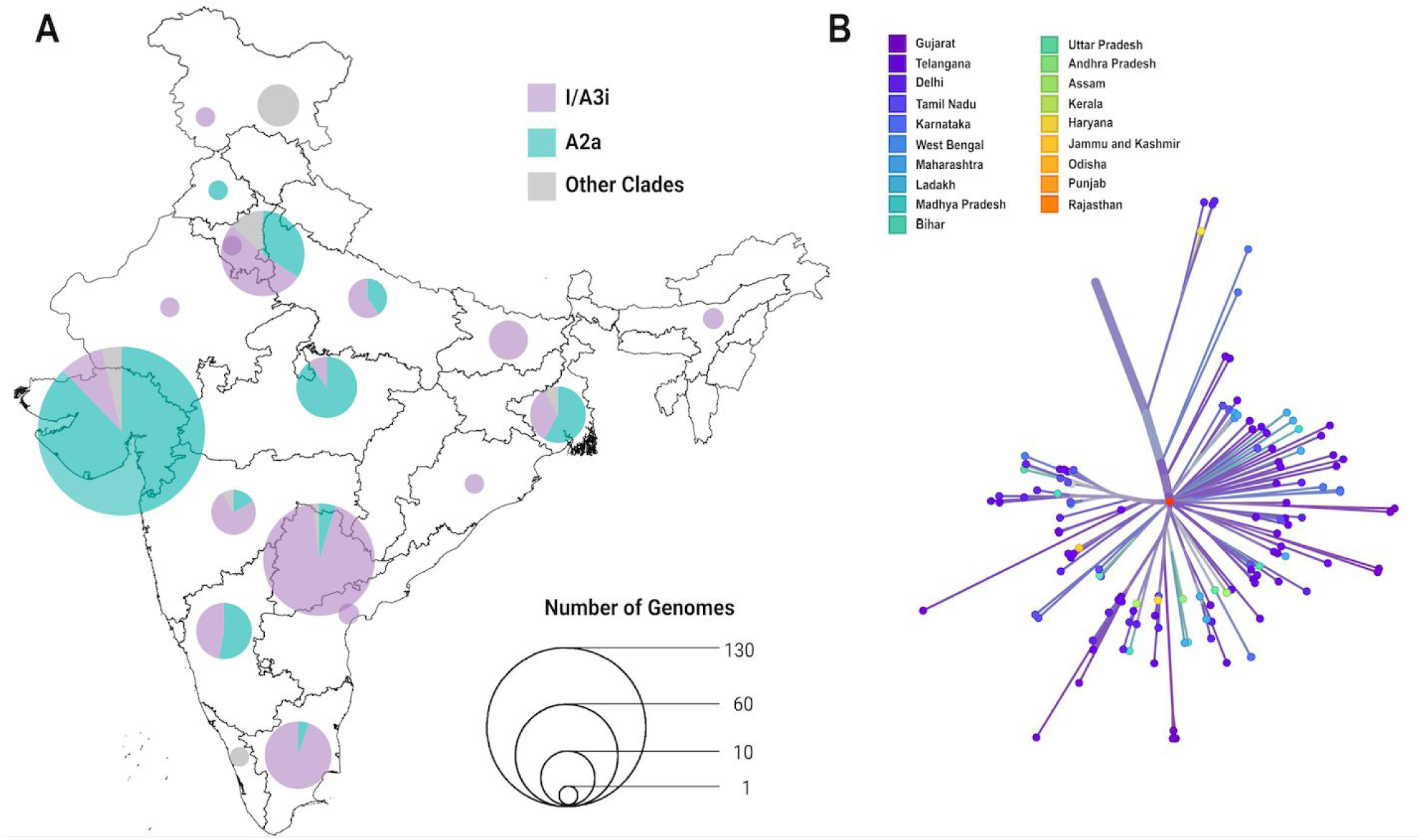
**A)** Proportion of the I/A3i clade (purple) and A2a (teal) in the genomes sequenced from different states of India. The proportion of the A2a clade (teal) is also shown for comparison, whereas all other clades are shaded grey. **B)** The short tree of the I/A3i clade diverging from a central point suggests a single point of introduction and spread across the different states.

Considering all the genomic data available from India, the I/A3i clade is represented in 145 genomes (41.2%) and represented from 16 of the 19 states from which the genomes originated. The geographical distribution and the proportion of the Clade I/A3i isolates are depicted in Figure 3. The states of Tamil Nadu, Telangana, Maharashtra, and Delhi have the highest proportions of this clade, followed by Bihar, Karnataka, Uttar Pradesh, West Bengal, Gujarat, and Madhya Pradesh. The states of Haryana, Jammu and Kashmir, Madhya Pradesh, Odisha, and Rajasthan have one genome each under Clade I/A3i **(Supplementary Data 6)**.

### Demographics of the patients

Of members in the Clade I/A3i, 43 were male (69.4%) while 19 were female (30.6%). The mean age was 36.8 years (CI 33.2 to 40.3 years). For A2a cluster, 72 were male (54.2%) and 61 were female (45.8%), while the mean age was 41.1 years (CI 37.9 to 44.2 years). While age was found to be significantly different between Clade I/A3i and other clades (p-value 0.000282), sex and clinical features were not found to be significantly different between Clade I/A3i and the other clades (p-value 0.13 and 0.72 respectively). The patient details for the Indian samples as provided by GISAID are available in **Supplementary data** 7.

## Discussion

Genomic evolution coupled with the appropriate tools like genome sequencing provides a unique opportunity to understand the spread and evolution of pathogens.^30,31^ The recent emergence of COVID-19 as a global pandemic and the emergence of the Open Data for SARS-CoV-2 genomes from across the globe facilitated by genomic databases like GenBank and GISAID has truly opened up new opportunities to understand the pathogen and its spread and evolution at an unprecedented rate.^4,32^ Whole-genome sequencing of SARS-CoV-2 has also been extensively used in understanding epidemics at a macro-level as well as at micro-levels, at hospitals.^33^

In this report, we describe a distinct cluster of sequences from genomes of SARS-CoV-2 sequenced and deposited from multiple laboratories across India, which we classify as the I/A3i clade. This distinct cluster could not be classified into any of the 10 clade annotations as described by Nextstrain, and was characterized by a unique combination of four variants which was shared by over 92% of the isolates falling in the cluster. The cluster was predominantly found in genomes from India; though additional members could also be found from genomes deposited in other countries, they form a minor proportion of the genomes from the respective countries. The Indian genomes constituted ∼63% of the global genomes for this cluster.

In-depth analysis of the genome cluster suggests a comparable rate of nucleotide substitutions with other predominant clades, though a gene-wise estimate of substitution suggests a distinct mode of evolution, driven by the Nucleocapsid (N) and Envelope (E) genes, and sparing of the Spike (S) gene in contrast to predominant diversity in the Spike (S) and Membrane protein (M) genes in A2a clade, the globally predominant clade.^34^ However, it has not escaped our attention that host genetic factors could modulate the evolution of the virus genome, and without large-scale host-genomic studies, the causal relationships cannot be conclusively established.

The cluster suggests a potential single introduction in February, followed by a country-wide spread, mostly affecting the South Indian states as evidenced by the tMRCA as well as the short cluster. Our analysis suggests that the Clade I/A3i was represented in almost all states from which genomes are available, barring a few. Members of the Clade I/A3i formed the predominant class of isolates from the states of Tamil Nadu, Telangana, Maharashtra, and Delhi and the second largest in membership in Bihar, Karnataka, Uttar Pradesh, West Bengal, Gujarat, and Madhya Pradesh.

Put together, the cluster of genomes (Clade I/A3i) forms a distinct cluster, predominantly found amongst Indian SARS-CoV-2 genomes, with limited representation outside the region. To the best of our knowledge, this is the first comprehensive study characterizing the distinct and predominant cluster of SARS-CoV-2 in India. This report also exemplifies the fact that timely and open access to genomic data can provide unique insights into the genetic epidemiology of pathogens.

## Author contributions

RKM, DTS and VS conceptualized and designed the study. SB and PM processed the sequencing data. SB, PM, and PS performed sequence data analysis and visualization. BJ performed the analysis for phylogenetic clustering, molecular characterization and quantification of relatedness. SK, LZ, SS and NG conducted the experiments. DTS, VS and BJ prepared the manuscript with inputs from RKM. All authors read and approved the manuscript.

## Conflicts of Interest

Authors declare no conflict of interest.

## Acknowledgements

Authors acknowledge the GISAID database and the contributors of genomic data, without which this analysis was not possible. The COVID-19 volunteer team of CCMB is thanked for help in initial sample processing. The full acknowledgements of data and contributors are available in **Supplementary Data 2**. Authors acknowledge funding from the Council of Scientific and Industrial Research (CSIR India). BJ is a recipient of the Junior Research Fellowship from CSIR India. The funders had no role in the design of the experiments, preparation of the manuscript or decision to publish.

## Data availability

All sequences generated in this study have been submitted to GISAID. The accession names of the samples and associated metadata are outlined in **Supplementary Data 1**.

## Supplementary Data

**Supplementary Data 1** – Metadata for the genome sequences submitted by our group

**Supplementary Data 2** – List of Indian GISAID submissions used in this study, and acknowledgements of all contributing authors

**Supplementary Data 3** – List of sites masked before variant analysis

**Supplementary Data 4** – Prior values used in the analysis of nucleotide substitution rates using BEAST

**Supplementary Data 5** – List of high quality global GISAID submissions used in Nextstrain analysis

**Supplementary Data 6** – State wise proportions of genomes belonging to the I/A3i clade

**Supplementary Data 7** – Metadata of clinically relevant information associated with the genomes deposited in GISAID from India.

**Supplementary Data 8 – Online resource**

An online and updated resource for SARS-CoV-2 genomes from India, their clade assignments and distribution across the country is available at **http://clingen.igib.res.in/genepi/phylovis/**

## Notes

### Competing Interest Statement

The authors have declared no competing interest.

