## Supplementary material for "A distinct phylogenetic cluster of Indian SARS-CoV-2 isolates": Fig S1

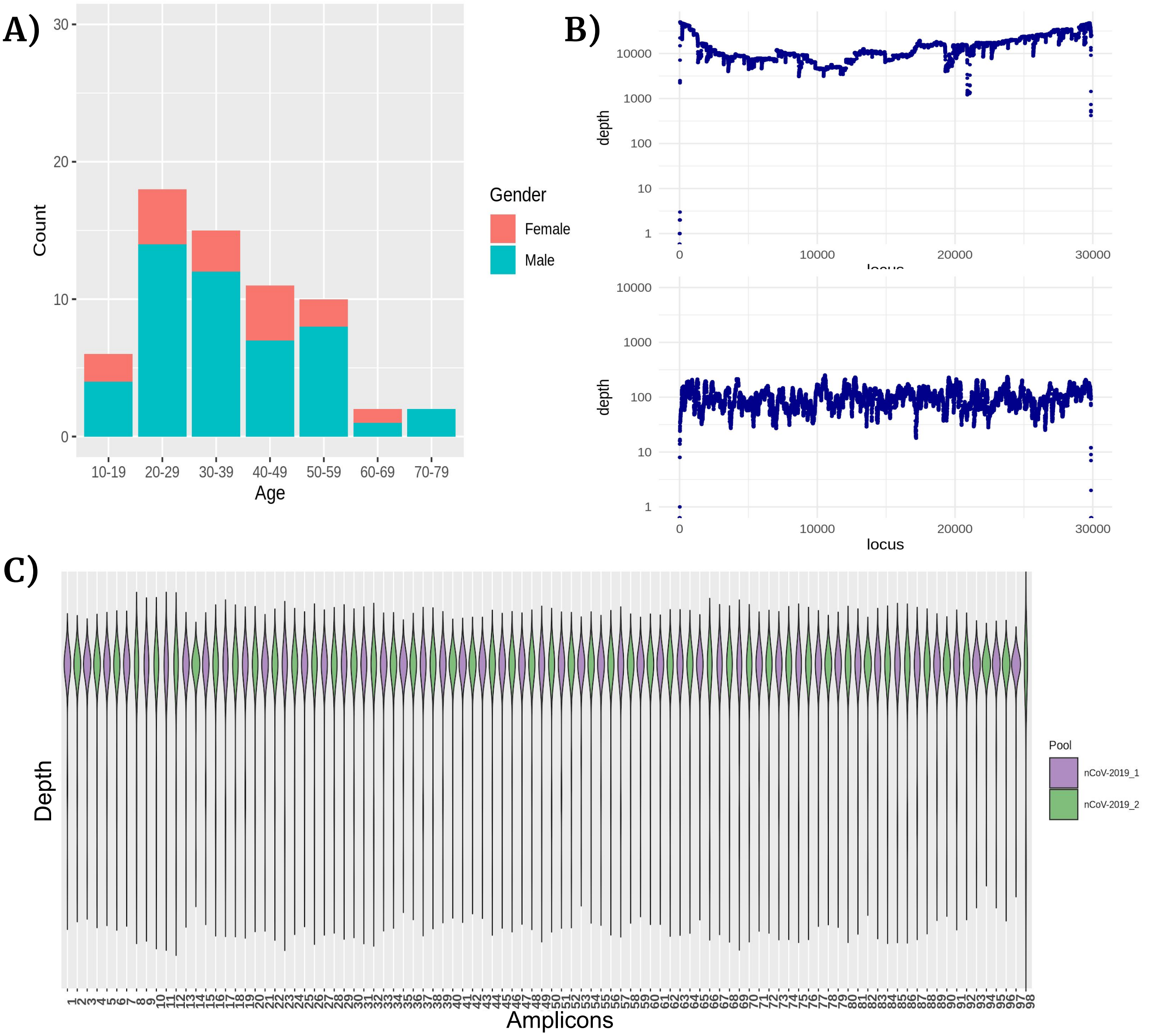


**Figure S1**: Profiles of SARS-CoV-2 sequences. **A)** Age and gender demographics of the sequenced individuals. **B)** A representative coverage plot for amplicon-based sequencing (top) and direct RNA sequencing (bottom). X-axis represents the 29.9 Kb genome and Y axis represents the depth in log scale. **C)** Average coverage obtained for each of the 98 amplicons in the ARTIC protocol. Except for the 98th amplicon (the last violin), all amplicons showed consistent amplification with many samples showing more than 10000x coverage.
